# An fMRI study of initiation and inhibition of manual responses in people who stutter

**DOI:** 10.1101/2020.09.07.286013

**Authors:** Charlotte E. E. Wiltshire, Jennifer Chesters, Saloni Krishnan, Máiréad P. Healy, Kate E. Watkins

**Author notes:** **Data Availability Statement:** The behavioural data that support the findings of this study are openly available on OSF at DOI 10.17605/OSF.IO/ZJSWA. The MRI data that support the findings of this study are openly available in Neurovault at https://identifiers.org/neurovault.collection:8669.

## Abstract

Developmental stuttering is a speech motor disorder characterised by difficulties initiating speech and frequent interruptions to the speech flow. Previous work suggests that people who stutter (PWS) have an overactive response suppression mechanism. Imaging studies of speech production in PWS consistently reveal greater activity of the right inferior frontal cortex, an area robustly implicated in inhibitory control of both manual and spoken responses. Here, we used a manual response version of the stop-signal task during fMRI to investigate neural differences related to response initiation and inhibition in PWS. Behaviourally, PWS were slower to respond to ‘go’ stimuli than people who are typically fluent (PWTF), but there was no difference in stop-signal reaction time. Our fMRI results were consistent with these behavioural results. The fMRI analysis revealed the expected networks associated with manual response initiation and inhibition in both groups. However, all contrasts between the two groups were characterised by overactivity in PWS relative to PWTF. This overactivity was significantly different for the initiation of responses (i.e. the ‘go’ trials) but not for response inhibition (i.e. the ‘stop’ trials). One explanation of these results is that PWS are consistently in a heightened inhibition state, i.e. areas of the inhibition network are more active, generally. This interpretation is consistent with predictions from the global response suppression hypothesis.

## Introduction

Contemporary evidence suggests developmental stuttering is a speech motor disorder with a distinct neural profile (Belyk, Kraft, & Brown, 2017; Brown, Ingham, Ingham, Laird, & Fox, 2005; Budde, Barron, & Fox, 2014; Neef, Anwander, & Friederici, 2015; Watkins, Smith, Davis, & Howell, 2008). Many brain areas that control speech production show differences, both functionally and structurally, in people who stutter (PWS) when compared with people who are typically fluent (PWTF). One such area is the right inferior frontal cortex, which is overactive in PWS during speech production; this overactivity is described as a ‘neural signature’ of developmental stuttering (Brown et al., 2005). The right inferior frontal cortex is also a key node in the network of brain areas involved in response inhibition (Aron, Robbins, & Poldrack, 2004; Chambers et al., 2007; Xue, Aron, & Poldrack, 2008). It is hypothesized that differences in inhibitory control may account, in part, for brain differences described in developmental stuttering (Alm, 2004; Hartwigsen, Neef, Camilleri, Margulies, & Eickhoff, 2019; Metzger et al., 2018; Neef et al., 2018, 2015; Xue et al., 2008). Here, a classic inhibition task was used with PWS whilst recording functional MRI images to test whether there are differences in the neural control of inhibition.

Hyperactivation in the frontal regions of the right hemisphere is robustly found in imaging work with PWS (Brown et al., 2005; Neef et al., 2015). One study found that this activation was positively correlated with stuttering severity (Neef et al., 2018) whilst another found an inverse correlation (Kell et al., 2009). In addition, evidence suggests that right hemisphere overactivity is abolished after therapy (De Nil, Kroll, Lafaille, & Houle, 2003; Kell et al., 2009), or during fluency enhancing techniques, such as choral reading (Fox et al., 1996). However, the mechanisms underlying this activity remain unclear. For example, it is unclear whether functional differences reflect general *traits* of developmental stuttering or specifically relate to moments of stuttering (*state* level). Recent evidence suggests that hyperactivation is more associated with state level stuttering: during dysfluent states relative to fluent ones, there was greater activation of inferior frontal and premotor cortex extending into the frontal operculum, bilaterally (Connally et al., 2018). Similarly, it is also unclear whether right frontal overactivation is related to error detection as a result of a stuttered moment or reflects an overactive inhibition signal. The DIVA (Directions Into Velocities of Articulators) model supports the idea that right frontal overactivity relates to error detection, considering the right posterior inferior frontal gyrus (IFG)/ventral pre-motor areas as a feedback control map (Tourville & Guenther, 2010), possibly detecting sensory-motor speech errors (Tourville, Reilly, & Guenther, 2008). Evidence that the right IFG overactivity reflects overactive inhibition, comes from a study that attempted to isolate the time course of right IFG activity during speech, from initiation to inhibition of the utterance (Neef et al., 2016). Compared with other regions activated by speaking, including the left IFG and temporal areas, the right IFG showed delayed peak activations, corresponding to the end of utterances. This temporal delay was observed in both PWS and PWTF but the peaks were larger in the right than left hemisphere in PWS. In addition, hyperactivity of right IFG correlated positively with stuttering severity, suggesting that it contributes to stuttering (Neef et al., 2018). Combined, these results support a ‘global suppression hypothesis’, which proposes that increased activation in right IFG represents in an increased tendency to globally (i.e. broadly, and non-specifically) inhibit motor responses, which contributes to stuttering.

The right IFG is part of a cortico-subcortical network of areas that controls movement initiation and inhibition (Aron & Poldrack, 2006; Aron et al., 2004). The hyperdirect pathway through the basal ganglia involves direct input from the IFG to ventral subthalamic nucleus (STN) bypassing the striatum (Chen et al., 2020). As the excitatory input from the cortex increases STN firing, which in turn excites the inhibitory output of the internal segment of the globus pallidus, the hyperdirect pathway is thought to provide rapid inhibition of basal ganglia output to the motor cortex (Nambu, Tokuno, & Takada, 2002). In vivo studies of the human brain using diffusion imaging identified white matter tracts connecting the STN with the supplementary motor complex (pre-SMA/SMA) and right IFG (Aron, Behrens, Smith, Frank, & Poldrack, 2007), which are connected in turn via the frontal aslant tract. This circuit may act to provide cortical control over this inhibitory pathway (Aron, Fletcher, Bullmore, Sahakian, & Robbins, 2003; Chambers et al., 2007). The involvement of the right IFG in stopping movement is supported by functional imaging results indicating that activity in the right IFG and the STN region is associated with inter-individual differences in movement inhibition (Aron & Poldrack, 2006); specifically, greater activity was associated with faster stopping times.

Recording and imaging subcortical structures to the level of detail described above is not feasible in PWS. Based on what is known from invasive methodologies in other populations, it is theorised that the output from the thalamus fails to provide appropriate timing cues for the initiation of speech movements to the motor networks, including SMA, premotor/motor cortex and cerebellum (Alm, 2004; Aron et al., 2004; Mink, 1996). Further evidence in support of a role for the basal ganglia or dopamine (or both) in developmental stuttering comes from pharmaceutical and lesion studies. Dopamine antagonists appear to improve fluency (Lavid et al., 1999; Maguire et al., 2000), whereas agonists worsen fluency (Anderson et al., 1999). In addition, increased levels of dopaminergic activity were described in the network of areas implicated in stuttering, including the medial prefrontal cortex, orbitofrontal cortex, insular cortex, auditory cortex as well as the ventral limbic cortical regions (Wu et al., 1997). Patients with neurogenic (acquired) stuttering have lesions encompassing the putamen (striatum), pallidum, cortical motor areas (Heuer, Sataloff, Mandel, & Travers, 1996), and the thalamus (Van Borsel, Van Der Made, & Santens, 2003). Whilst such lesion studies suggest a link between the basal ganglia-cortical circuits and stuttering, the size of the lesions makes it difficult to establish a causal relationship between damage to a specific region and behaviour (Alm, 2004). A key piece of evidence that is missing is whether PWS have hyperactivity in this network during an inhibition task, independent of speech and stuttering.

The stop-signal task paradigm has been used in conjunction with fMRI to robustly isolate inhibition responses (Aron & Poldrack, 2006; Chevrier, Noseworthy, & Schachar, 2007; Ray Li, Yan, Sinha, & Lee, 2008; Xue et al., 2008). During this task, participants produce a button-press response to a visual stimulus as quickly as possible. On a small percentage of randomly inserted trials, participants are cued to inhibit their response by an auditory tone (the “stop signal”). The timing of the stop signal is adjusted for each individual to determine the interval needed such that participants fail to inhibit their action 50% of the time and allow estimation of a stop-signal reaction time (SSRT; the time needed to successfully inhibit a response). This paradigm was adapted to measure inhibition of speech responses and used to demonstrate that both motor and spoken response inhibition evoke common neural activity in the right IFG (Xue et al., 2008). This suggests that inhibitory control is a domain general process.

Previously, a behavioural version of the stop-signal task was used to test manual response inhibition in adults who stutter and showed that PWS have a longer SSRTs compared with PWTF (Markett et al., 2016). Similarly, use of the GO/NOGO task also showed that children who stutter had a decreased ability to inhibit manual responses compared with matched controls (Eggers, De Nil, & Van Den Bergh, 2013). These findings suggest that PWS have problems enacting an inhibitory response, which is contrary to the prediction from the global suppression hypothesis that PWS have a greater tendency to inhibit motor responses (Neef et al., 2018). It is clear, therefore, that the addition of fMRI recording during the stop-signal task could help to further understand the neural basis of response inhibition in PWS.

Limited evidence exists directly linking stuttering with neural inhibition. One study recorded functional MRI whilst PWS performed a manual GO/NOGO task (Metzger et al., 2018). This study found increased activity in the basal ganglia and thalamus and particularly in the substantia nigra during response preparation. Importantly, task-related activity in the substantia nigra correlated positively with the trait of stuttering severity. In addition, the globus pallidus and the thalamus showed increased network synchronization with the IFG in PWS compared with PWTF (Metzger et al., 2018). Similarly, other work shows a positive correlation between white matter connectivity of the thalamus with right IFG and stuttering severity (Neef et al., 2018). Together, these findings provide evidence for abnormal function of the basal-ganglia-thalamo-cortical network involved in inhibition of manual responses in PWS. These findings in the manual domain indicate differences in inhibitory control beyond speech motor control, which would be consistent with the idea that speech and non-speech movements share an inhibitory control network (Xue et al., 2008).

Here, we used fMRI during a stop-signal task to capture manual initiation and inhibition responses of PWS and PWTF. Our aim was to investigate neural differences related to response initiation and inhibition in PWS using the stop-signal paradigm. According to the global suppression hypothesis, shorter SSRT behaviourally and hyperactivation of the right hemisphere inhibition network during ‘go’ and ‘stop’ trials were predicted. Using the manual version of the stop-signal task allowed us to address whether differences in inhibitory control extend outside the speech domain and might be considered domain general.

## Pre-scan behavioural task

### Participants

38 adult PWS and 21 matched controls participated in the study. Groups were balanced for gender, age, years of education and ethnicity (see Table 1, below). In the stuttering group, stuttering severity ranged from moderate to very severe as measured by the SSI (Stuttering Severity Instrument-4; Riley 2009) and participants had no diagnosis of other speech or language disorders. Participants had not undergone speech therapy for at least 6 months prior to testing. They had corrected or corrected-to-normal vision and hearing.

**Table 1.**
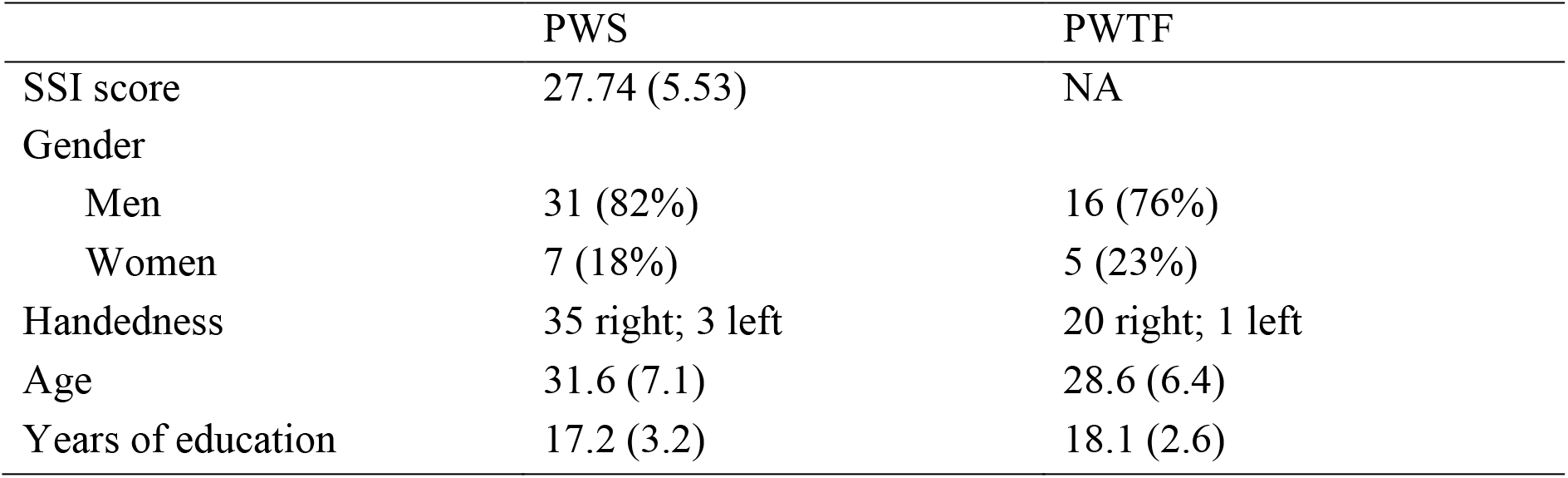
Demographics of participants. Group counts and proportions (%) of group are shown for gender. The means and SD are provided for SSI score, age and years of education.

The University of Oxford Central University Research Ethics Committee approved the study. Participants gave informed written consent to participate in the study, in accordance with the Declaration of Helsinki, and with the procedure approved by the committee.

### Pre-scan task

The stop-signal task was run before the scan session in order to determine the stop-signal delay (SSD) to be used during scanning for each participant. The task was identical to the pre-scan task described in Xue et al., (2008). 240 Go trials and 80 Stop trials were used to estimate the SSD for manual responses. On Go trials, each trial started with a white fixation cross, presented on a grey background for 500 ms. After which, a visual stimulus appeared on the screen for one second. Participants were presented with a left or right facing arrow (< or >). Participants responded by pressing one of two buttons with the right index finger corresponding to the left/right direction of the arrow. The inter-trial delay was two seconds. Stop trials were visually identical to go trials except that an auditory cue (500ms) was played at an interval after presentation of the visual stimulus (arrow). Participants were instructed to respond as fast as possible to the visual stimulus and were told that it would not always be possible to stop in response to the auditory cue. The delay was changed adaptively according to the subject’s behaviour. If the participant inhibited successfully on a stop trial, then inhibition was made less likely on the next stop trial by increasing the SSD by 50ms, thus increasing the time that the visual stimulus was on the screen before the stop signal and the likelihood that the participant would begin to respond before the auditory stop signal was presented (i.e. making the stopping more difficult). If the participant failed to inhibit, the SSD was decreased by 50ms, thus giving participants less time to initiate a Go response before the cue was played and the response was more easily stopped or inhibited. The SSD was varied in this way using a staircase algorithm. An individual participant’s SSD was computed as the SSD at which the probability of a participant successfully inhibiting on a Stop trial was 50%. This SSD was used during scanning, thereby ensuring that the task was equally difficult for each participant. As SSD was varied to yield a 50% chance of stopping, the stop-signal reaction time (SSRT) was estimable by subtracting the mean SSD from the median reaction time (RT) of correct Go responses. The SSRT provides a measure of the speed of the stopping process: a short SSRT indicates a quick stopping process and a long SSRT indicates a slow stopping process.

### Behavioural results

The results of the behavioural study are summarised in table 2 and visualised in figure 1.

**Table 2.** Group performance on the stop-signal behavioural task.

|  | PWS | PWTF |
| --- | --- | --- |
| Go reaction time | 542.41 (92.18) | 488.61 (76.06) |
| SSD | 325.23 (122.91) | 277.56 (98.97) |
| SSRT | 217.18 (64.92) | 211.05 (67.22) |
| Percent responding on go trials | 96.24 (5.02) | 98.91 (0.96) |
| *Group means (standard deviation) are shown.* |  |  |

Independent t-tests run in R using the base R ‘stats’ package (R Core Team, 2019) indicated that, for the Go trials reaction time, PWS responded significantly more slowly than PWTF ((t(57) = 2.28, *p* = 0.027, d = 0.64). PWS and PWTF showed no difference in SSD (t(57) = 1.53, *p* = 0.13) nor SSRT (t(57)=0.34, *p* = 0.73).

**Figure 1.**
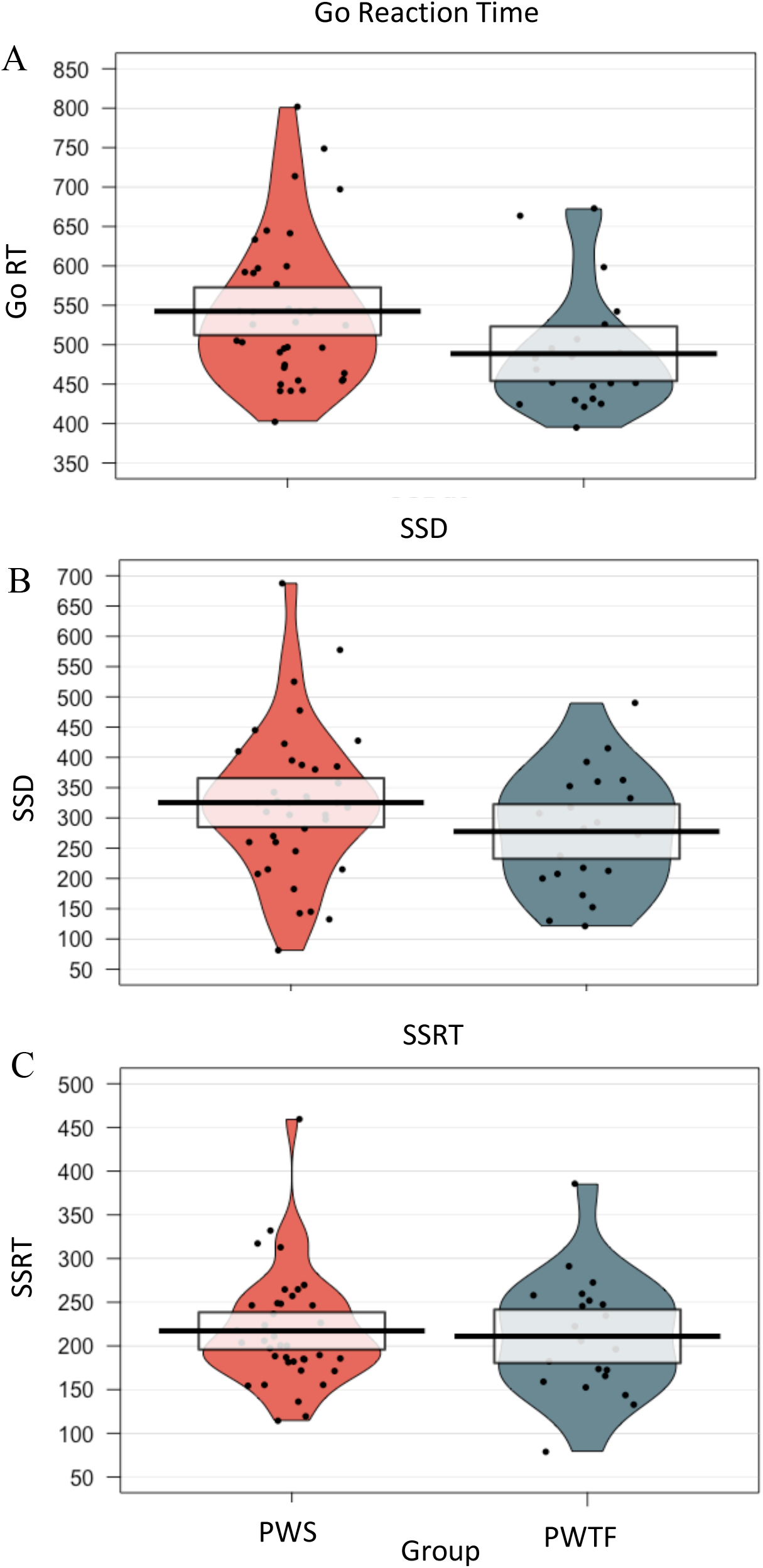
Group performance on the behavioural stop-signal task. Violin plots are shown for the PWS (orange) and PWTF (blue) groups for A) Go Reaction Time; B) Stop Signal Delay; C) Stop Signal Reaction Time. Data points for individual subjects are shown by the small black dots. The group mean is indicated by the solid black line and the 95% confidence interval by the white shaded box. The colour shading indicates the distribution of data for each group.

## Task fMRI

### Participants

The SSD from the behavioural task was used during scanning to approximate the delay at which participants would successfully stop 50% of the time and fail to stop 50% during stop trials. This approximation did not always result in a near-equal distribution of stopping responses during scanning. Therefore, data were included in the following MRI analysis for participants who produced at least 10/24 successful or unsuccessful stops during the scan. This resulted in exclusion of data from two participants (both PWS). In addition, data from seven participants (five PWS and two PWTF) were excluded for technical reasons (e.g. button box failure, scanner issues) and a further three participants (one PWS and two PWTF) were excluded for excessive movement during scanning (>2mm average absolute movement).

After these exclusions, data from 30 PWS (6 women, mean age = 31.6, SD = 7.4) and 17 PWTF (5 women, mean age = 28.5, SD = 6.0) were analysed.

### fMRI SSRT task

The stop-signal paradigm was used to assess manual responses as described in Xue et al., (2008). The task comprised 144 Go and 48 Stop trials. The paradigm in the scanner was the same as described above, except that the staircasing algorithm was replaced with a fixed, individualised, SSD, which estimated the probability of successfully inhibiting on 50% of trials.

### MRI data acquisition

Image data were acquired using a 3T Siemens magnetom Prisma scanner using a 64-channel head and neck coil. Echo-planar images were acquired with 72, 2-mm isometric slices with multiband acceleration factor of 8 (TR/TE = 720/33, Flip Angle = 53 deg, FOV = 208, in-plane resolution of 2 × 2 mm). High-resolution T1-weighted structural images of the whole brain were also acquired with an MPRAGE protocol (PAT2, 1mm isotropic, TR/TE = 2400/3.98). Audio stimuli (i.e. the stop signal) were delivered via OptoActive active noise-cancelling headphones (Optoacoustics ltd, Israel).

### Imaging preprocessing and statistics

Data for each participant were preprocessed using standard parameters in FMRIB Software Library (FSL version 6.0.1 http://www.fmrib.ox.ac.uk/fsl, [Jenkinson, Beckmann, Behrens, Woolrich, & Smith, 2012]). For each participant, the whole-brain T1-weighted image was skull stripped using the Brain Extraction Tool (BET; part of FSL). Functional data were processed at the subject-level using FMRI expert analyses tool (FEAT, v 6.0). A temporal high-pass filter with a cut-off of 90s was used to remove low-frequency fluctuations in the signal. Standard motion correction was applied (MCFLIRT). Data were smoothed with a 5-mm full-width-at-half-maximum Gaussian smoothing kernel. *B*0 unwarping was conducted using the fieldmap images and PRELUDE and FUGUE software running in FSL (Jenkinson et al., 2012). All fMRI volumes were first registered to a reference image (increased SNR and contrast but with same distortions) and then aligned to the individual’s structural scan using brain boundary registration (BBR), implemented using FMRIB’s Linear Image Registration Tool (FLIRT). They were then registered to 2-mm MNI standard space using FMRIB’s Nonlinear Image Registration Tool (FNIRT) for group analyses.

Group comparisons were implemented using FMRIB’s Local Analysis of Mixed Effects stage 1 (Woolrich et al., 2004). For group averages, results are reported using a cluster-forming threshold Z > 3.1, and extent-threshold of p < .05 corrected. To protect against possible false negative errors in the between-group contrasts, we used a reduced cluster-forming threshold of Z > 2.3 and extent threshold of p < .05 (corrected).

### fMRI results

#### Go trials

The controls showed focal activation of the left pre- and post-central gyri at the level of the hand representation, the left putamen extending to the opercular cortex, the SMA and extensively in the cerebellum bilaterally (see Fig. 2 and Table 3). This pattern of activation was consistent with the task which involved a simple button press with the right index finger in response to the visually presented arrow. There was also activity bilaterally in the occipital cortex evoked by the visual stimulus.

**Figure 2.**
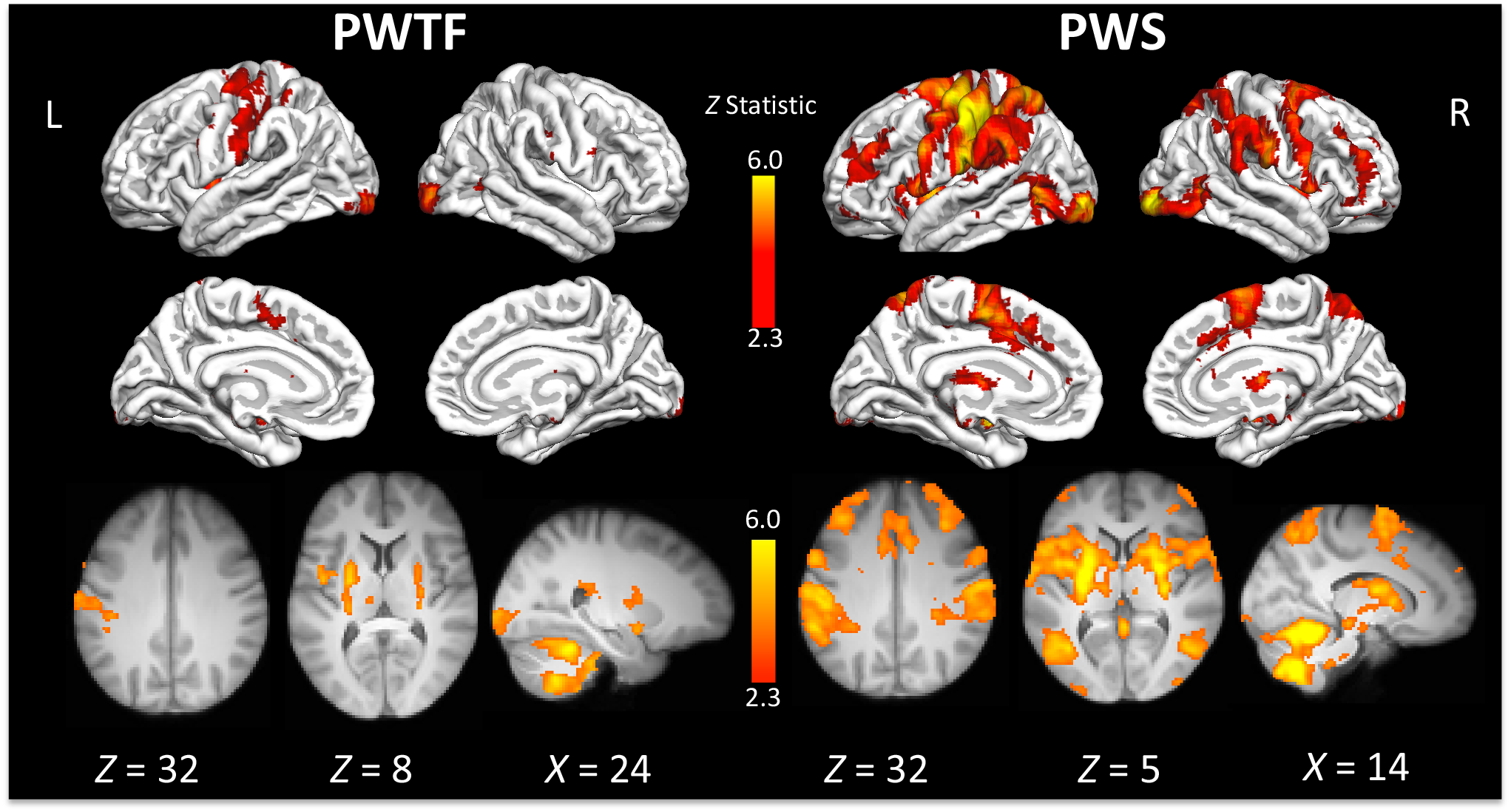
Activity during ‘go’ trials for PWTF and PWS. Coloured areas indicate statistical maps (thresholded at Z>2.3 for visualisation purposes) overlaid on the cortical surface using FreeSurfer or on slices through the brain volume at the coordinate indicated below each image. PWTF = people who are typically fluent. PWS = people who stutter. L – left; R – right. See Tables 3 and 4 for a list of areas significantly activated for each group.

**Table 3.**
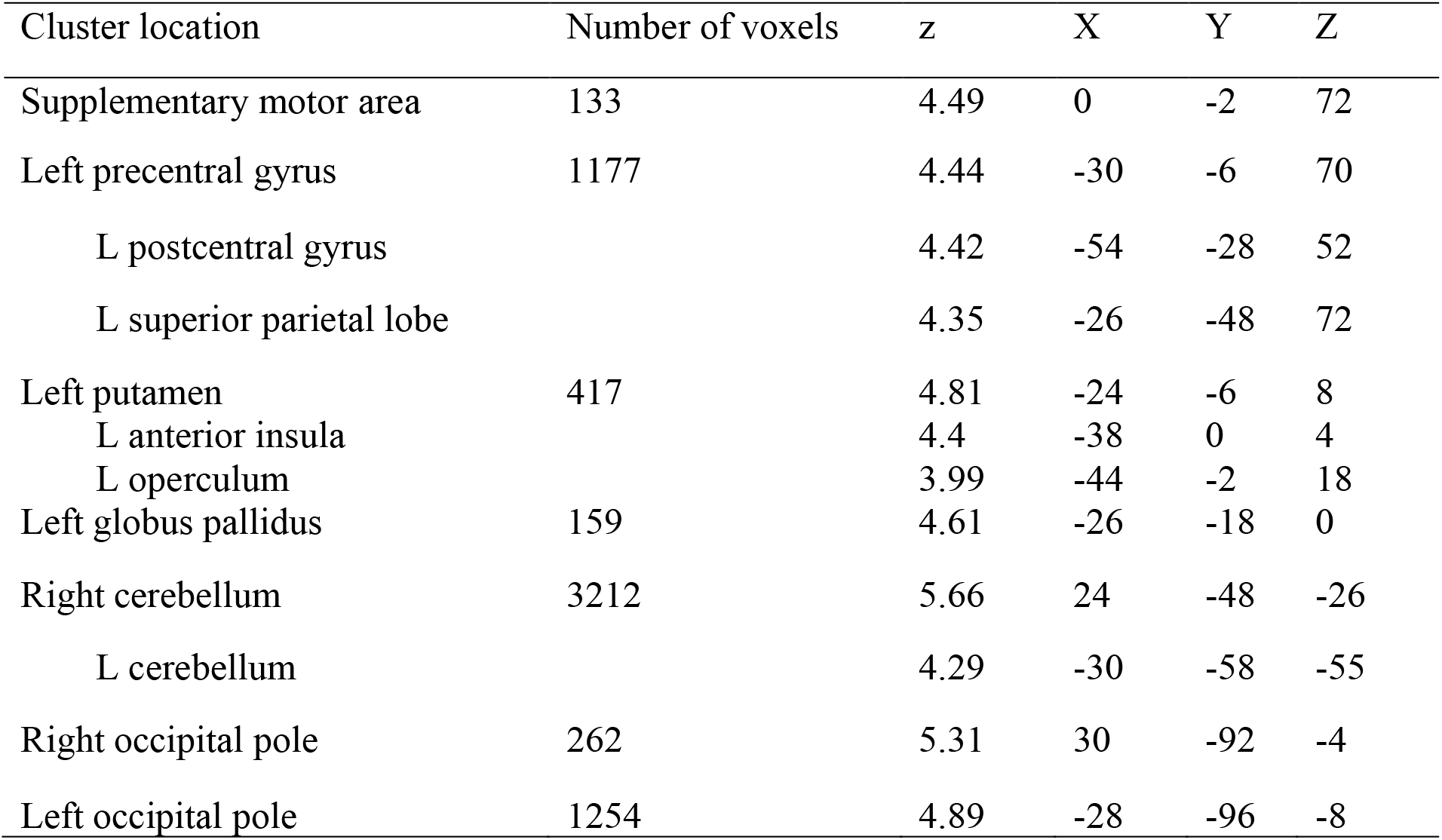
Activation peaks and coordinates for peaks in clusters significantly activated in PWTF during ‘go’ trials. Cluster forming threshold Z > 3.1, extent threshold p < .05 corrected. The cluster size, peak Z statistic, and MNI coordinates of selected peaks are provided.

In contrast to the focal (and expected) pattern of activity in the controls, PWS had extensive and widespread activation of the frontal operculum, precentral gyrus, SMA, putamen, and cerebellum, all bilaterally (see Fig. 2 and Table 4). The left postcentral gyrus and supramarginal gyrus bilaterally were also robustly activated. There was also activity in the anterior portion of the middle frontal gyrus bilaterally. Visual cortex activity extended from the pole to include the lateral occipital cortex bilaterally.

**Table 4.** Activation peaks and coordinates for peaks in clusters significantly activated in PWS during ‘go’ trials. See Table 3 for details.

| Cluster location | Number of voxels | z | X | Y | Z |
| --- | --- | --- | --- | --- | --- |
| Right middle frontal gyrus | 636 | 4.63 | 44 | 44 | 24 |
| Left middle frontal gyrus | 691 | 4.46 | -38 | 34 | 32 |
| Cortical and subcortical motor areas | 22195 |  |  |  |  |
| R putamen |  | 6.89 | 26 | -2 | 4 |
| Supplementary motor area |  | 6.09 | 0 | -4 | 46 |
| L putamen |  | 7.57 | -26 | -6 | 6 |
| L precentral gyrus |  | 7.09 | -50 | -18 | 56 |
| L postcentral gyrus |  | 6.86 | -42 | -26 | 48 |
| Right postcentral gyrus | 3143 | 5.2 | 34 | -40 | 36 |
| Cerebellum and occipital lobe | 13938 | 7.91 | 14 | -52 | -20 |
| L cerebellum |  | 6.32 | -31 | -49 | -28 |
| R cerebellum |  | 7.34 | 18 | -62 | -48 |
| R occipital pole |  | 7.41 | 32 | -92 | -4 |

The contrast between groups confirmed that this pattern of widespread greater activity in PWS was statistically significant. PWS had significantly greater activity relative to PWTF in the inferior frontal gyrus, caudate nucleus and putamen bilaterally, and in the left precentral cortex and parietal operculum (see Fig. 3).

**Figure 3.**
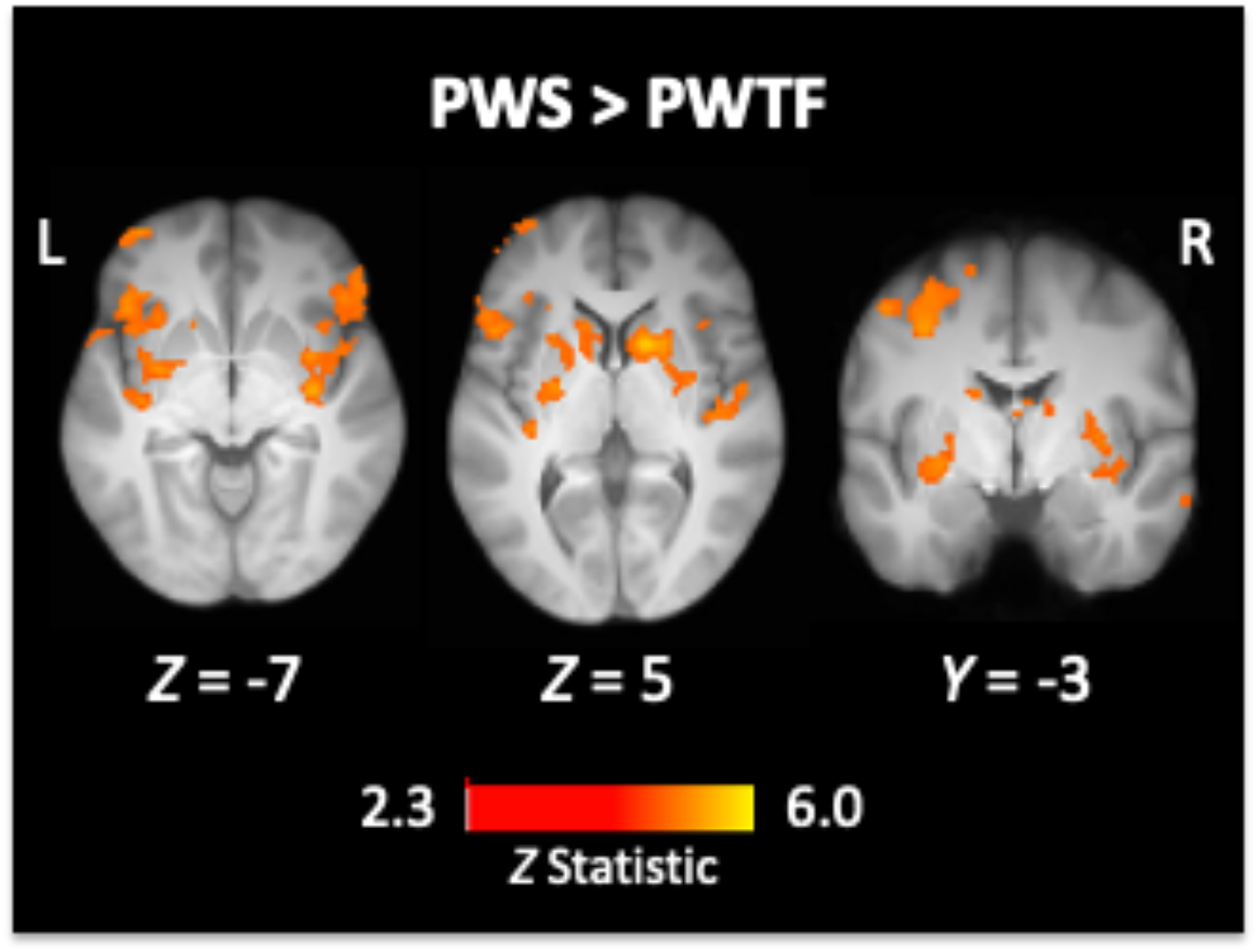
Areas with significantly greater activity in PWS than PWTF during ‘go’ trials. See legend to Figure 2 for details. See Table 5 for a list of areas significantly activated for this contrast.

**Table 5.** Activation peaks and coordinates for peaks in clusters significantly activated in PWS more than PWTF during ‘go’ trials. Cluster-forming threshold Z > 2.3, extent threshold p < .05 corrected. See Table 3 for details.

| Cluster Location | Number of voxels | z | X | Y | Z |
| --- | --- | --- | --- | --- | --- |
| Left inferior frontal gyrus | 1483 |  |  |  |  |
| L pars orbitalis |  | 3.96 | -44 | 26 | -10 |
| L pars opercularis |  | 3.53 | -50 | 16 | 2 |
| Right striatum and operculum | 1678 | 4.78 | 14 | 10 | 4 |
| R inferior frontal gyrus (triangularis) |  | 3.47 | 52 | 24 | -2 |
| R anterior insula |  | 3.69 | 34 | 6 | -14 |
| R caudate nucleus<sup>†‡</sup> |  | 3.5 | 14 | 4 | 16 |
| R putamen |  | 4.2 | 34 | -12 | -4 |
| Left striatum | 698 |  |  |  |  |
| L putamen |  | 3.77 | -22 | 0 | -4 |
| L caudate nucleus<sup>‡</sup> |  | 3.34 | -14 | 6 | 14 |
| Left precentral gyrus<sup>†</sup> | 831 | 4 | -36 | -2 | 44 |
| Left parietal operculum | 484 | 3.64 | -60 | -24 | 24 |
| Right medial parietal cortex | 397 | 4.18 | 12 | -66 | 56 |

<sup>†</sup>survived cluster thresholding at $z > 3.1$ for the group contrast
<sup>‡</sup>symmetric peaks in the left and right caudate nucleus

#### Successful stop trials

On the ‘successful stop’ trials, PWTF activated the frontal operculum extending to IFG, anterior insula and putamen, middle frontal gyrus, postcentral gyrus, supramarginal gyrus, and cerebellum bilaterally, and the SMA extending to the cingulate motor area. As for the ‘go’ trials, the pattern of activation in PWS for the successful stop trials was extensive and looked like an amplified version of the network seen in PWTF. When the two groups were contrasted, this pattern of overactivity in PWS was not significantly different to that in PWTF.

**Table 6.** Activation peaks and coordinates for peaks in clusters significantly activated in PWTF during ‘successful stop’ trials. See Table 3 for details.

| Cluster Location | Number of voxels | z | X | Y | Z |
| --- | --- | --- | --- | --- | --- |
| Right middle frontal gyrus | 429 | 4.51 | 40 | 40 | 30 |
| Left middle frontal gyrus | 305 | 4.68 | -38 | 30 | 38 |
| Right striatum and operculum | 1603 |  |  |  |  |
| R frontal operculum |  | 5.05 | 42 | 14 | -2 |
| R inferior frontal gyrus (opercularis) |  | 4.93 | 56 | 12 | 0 |
| R anterior insula |  | 5.63 | 42 | 8 | -2 |
| R putamen |  | 5.25 | 26 | 0 | 10 |
| Left striatum and operculum | 2444 |  |  |  |  |
| L frontal operculum |  | 5.19 | -32 | 16 | 14 |
| L inferior frontal gyrus (opercularis) |  | 4.41 | -48 | 12 | 11 |
| L anterior insula |  | 5.34 | -40 | 2 | 4 |
| L putamen |  | 4.95 | -28 | -16 | 5 |
| Right dorsal motor cortex | 1283 |  |  |  |  |
| R cingulate motor area |  | 3.92 | 2 | 13 | 41 |
| R precentral gyrus |  | 5.09 | 36 | -4 | 64 |
| R SMA |  | 4.88 | 2 | -6 | 74 |
| Left sensorimotor cortex | 5545 |  |  |  |  |
| L precentral gyrus |  | 5.58 | -30 | -6 | 70 |
| L postcentral gyrus |  | 5.17 | -42 | -28 | 42 |
| L supramarginal gyrus |  | 5.4 | -60 | -36 | 22 |
| Right inferior parietal | 3170 |  |  |  |  |
| R postcentral gyrus |  | 4.55 | 62 | -14 | 36 |
| R supramarginal gyrus |  | 5.2 | 64 | -36 | 36 |
| Cerebellum | 7174 |  |  |  |  |
| R cerebellum |  | 6.09 | 26 | -48 | -26 |
| L cerebellum |  | 5.66 | -46 | -61 | -28 |
| Right occipital pole | 333 | 5.5 | 30 | -92 | -4 |

**Table 7.** Activation peaks for PWS during successful stop trials. Using Z>3.1 resulted in clusters of very large extent spanning multiple anatomical areas. We report locations for these clusters using Z>4.3 cluster-forming threshold for clarity (labelling truncated for clusters with < 50 voxels in extent). The cluster size, peak Z statistic, and MNI coordinates of selected peaks are provided.

| Cluster Location | Number of voxels | z | X | Y | Z |
| --- | --- | --- | --- | --- | --- |
| Right middle frontal gyrus | 1045 | 6.01 | 42 | 48 | 24 |
| Right frontal pole | 118 | 5.78 | 22 | 44 | -12 |
| Left middle frontal gyrus | 435 | 6.28 | -40 | 36 | 34 |
| Right striatum and operculum | 2461 |  |  |  |  |
| R inferior frontal gyrus |  | 5.84 | 57 | 15 | -2 |
| R insula |  | 7.15 | 46 | 14 | -6 |
| R putamen |  | 7.12 | 28 | -2 | 8 |
| Left striatum and operculum | 2518 |  |  |  |  |
| L operculum/insula |  | 6.61 | -46 | 14 | -6 |
| L inferior frontal gyrus |  | 6.47 | -52 | 12 | 2 |
| L putamen |  | 7.35 | -24 | 2 | 6 |
| Left medial and lateral cortex | 8958 |  |  |  |  |
| L cingulate gyrus |  | 5.2 | -5 | 5 | 42 |
| L SMA |  | 6.1 | -2 | -4 | 55 |
| L operculum |  | 7.3 | -64 | -22 | 16 |
| L postcentral gyrus |  | 7.39 | -42 | -28 | 48 |
| L supramarginal gyrus |  | 6.7 | -61 | -47 | 36 |
| Right precentral gyrus | 108 | 4.96 | 62 | 8 | 26 |
| Left precentral gyrus | 93 | 5.41 | -58 | 6 | 30 |
| Cerebellum | 6342 |  |  |  |  |
| R cerebellum |  | 7.9 | 32 | -48 | -48 |
| L cerebellum |  | 7.19 | -32 | -58 | -12 |
| Right supramarginal gyrus | 4371 | 6.62 | 64 | -36 | 30 |
| R postcentral gyrus |  | 5.99 | 59 | -16 | 30 |
| Right superior parietal lobe | 63 | 5.35 | 12 | -54 | 56 |
| Left lateral occipital cortex | 431 | 6.04 | -52 | -68 | 2 |
| Right occipital pole | 369 | 7.12 | 32 | -92 | -4 |
| Left occipital pole | 474 | 6.39 | -26 | -100 | -6 |

#### Unsuccessful stop trials

For the ‘unsuccessful stop’ trials, PWTF showed activity in the left postcentral gyrus, putamen and thalamus, and bilaterally in the supramarginal gyrus, SMA extending to cingulate motor cortex, frontal opercular cortex, and cerebellum. In PWS, a similar but more robust pattern of activation was seen. These two patterns of activation were not significantly different, however.

**Table 8.** Activation peaks and coordinates for peaks in clusters significantly activated in PWTF during ‘unsuccessful stop’ trials. See Table 3 for details.

| Cluster Index | Number of voxels | z | X | Y | Z |
| --- | --- | --- | --- | --- | --- |
| Right middle frontal gyrus | 220 | 4.54 | 42 | 42 | 30 |
| Right frontal operculum | 741 | 5.15 | 50 | 14 | -4 |
| Left frontal operculum | 1253 | 5.34 | -46 | 12 | -6 |
| Medial frontal cortex | 1616 |  |  |  |  |
| L SMA |  | 5.38 | 2 | -6 | 74 |
| L Cingulate cortex |  | 4.18 | 2 | 17 | 40 |
| Left putamen | 182 | 4.45 | -28 | -22 | 2 |
| Left sensorimotor cortex | 3484 |  |  |  |  |
| L postcentral gyrus |  | 5.16 | -56 | -28 | 48 |
| L supramarginal gyrus |  | 5.13 | -57 | -45 | 26 |
| Right supramarginal gyrus | 1532 | 5.41 | 56 | -40 | 36 |
| Left cerebellum | 1776 | 4.95 | -38 | -50 | -32 |
| Right cerebellum | 3230 | 5.69 | 34 | -54 | -26 |

**Table 9.** Activation peaks and coordinates for peaks in clusters significantly activated in PWS during ‘unsuccessful stop’ trials. See Table 3 for details.

| Cluster Index | Number of voxels | z | X | Y | Z |
| --- | --- | --- | --- | --- | --- |
| Left middle frontal gyrus | 510 | 4.82 | -26 | 54 | 26 |
| Right middle frontal gyrus | 121 | 4.32 | 26 | 46 | -10 |
| Left middle frontal gyrus | 115 | 4.15 | -40 | 34 | 34 |
| Left frontal opercular cortex | 3319 | 7.38 | -46 | 14 | -4 |
| L putamen |  | 4.91 | -24 | -6 | 10 |
| Right frontal opercular cortex | 3839 | 6.71 | 54 | 12 | 8 |
| Right putamen | 202 | 4.66 | 22 | -2 | 10 |
| Left sensorimotor extending to medial frontal | 9979 | 7.52 | -60 | -46 | 34 |
| SMA |  | 4.92 | 0 | 20 | 61 |
| Cingulate cortex |  | 5.25 | 0 | 20 | 40 |
| L parietal operculum |  | 5.98 | -60 | -22 | 18 |
| L supramarginal gyrus |  | 7.04 | -62 | -46 | 30 |
| Right supramarginal gyrus | 4094 | 6.21 | 62 | -46 | 30 |
| Left cerebellum | 1956 | 5.71 | -34 | -58 | -56 |
| Right cerebellum | 4660 | 6.95 | 20 | -64 | -50 |

## Discussion

We tested whether there were differences in the neural control of the initiation and inhibition of a manual response in people who stutter (PWS) and people who are typically fluent (PWTF) in the context of the stop-signal paradigm. 38 PWS and 21 PWTF completed the behavioural version of the manual stop signal task (Xue et al., 2008). Of these participants, we also compared fMRI task data in 30 PWS and 17 PWTF. During the task, participants responded to a visual stimulus (left or right arrow) with their right index finger. On randomly inserted trials, participants heard an auditory cue, which indicated they should inhibit their response. Previous work suggests that PWS have an overactive response inhibition mechanism (Neef et al., 2018). According to the global suppression hypothesis, shorter SSRT and hyperactivation of the right hemisphere inhibition network were expected. Contrary to this prediction, in the current study, PWS had longer reaction times on ‘go’ trials than PWTF. There was no significant difference in the speed of the stopping process (SSRT). The patterns of fMRI activity during task were consistent with these behavioural results. During ‘go’ trials, both groups activated the expected motor control areas related to the task demands (a simple button press in response to a visual stimulus). In contrast to the focal pattern of activity in PWTF, however, PWS had extensive and widespread activation for performance of this simple task. PWS showed significantly more activation than PWTF in the inferior frontal gyrus, caudate nucleus and putamen bilaterally, and in the left precentral cortex and left parietal operculum. There was a similar pattern of activation during ‘successful stop’ trials, which also included activation of the supramarginal gyrus and cingulate cortex. As for the ‘go’ trials, the pattern of activation in PWS in the successful stop trials looked like an amplified version of that seen in PWTF, but these differences were not statistically significant.

Taken together, these results provide evidence for differences between PWS and PWTF in performance of a simple manual response and the related neural activity in the context of a task requiring inhibition. PWS show a pattern of overactivation for both ‘go’ and ‘stop’ trials that corresponds to the inhibitory control network (Aron et al., 2007, 2004).

### Behavioural results

A simple prediction arising from the global suppression hypothesis (Neef et al., 2018) is that PWS would have shorter stopping responses due to a constant heightened inhibition signal. Our results indicate that whilst PWS were significantly slower to initiate a response (go reaction time), the stopping response (SSRT) was not different to that in PWTF. This suggests that PWS do not differ in the time it takes to inhibit a manual response. The behavioural results of our study contrast with those from a previous study of PWS that also employed the manual stop signal task and found that PWS had longer stopping responses (SSRTs) than PWTF (Markett et al., 2016). Further study is required to resolve these discrepant findings. In the previous study (Markett et al., 2016), the effect size for the significant group difference of SSRT was d = 0.61 (n= 28 per group). To detect this moderate effect size, we would need 33 participants per group (assuming 80% power). Here, we had slightly below this total sample size with 38 PWS and 21 PWTF. We were therefore slightly underpowered to detect a similar effect size. Even so, the distributions of SSRTs for our two groups were very closely matched and there was no evidence of a trend towards longer SSRTs in the PWS relative to the PWTF.

The significantly longer reaction times for ‘go’ responses in PWS found in the current study were unexpected but could be explained in two ways. One explanation is that PWS have greater difficulty enacting a response under temporal uncertainty. For example, the previous study (Markett et al., 2016) used two tasks to estimate ‘go’ reaction times: one task had ‘go’ trials only and used fixed inter-trial intervals, providing strong temporal predictability for when a ‘go’ response was required. The other task involved trials with varied inter-trial-intervals, which provide less temporal predictability. PWS only showed longer reaction times relative to PWTF when the timing of the trials was unpredictable (Markett et al., 2016). The authors suggest that this difference may be due to problems relying on internally generated timing compared with the externally generated timing provided by the predictability of the fixed inter-trial-intervals (Alm, 2004; Markett et al., 2016). In the current study, a fixed inter-trial-interval was used, however ‘go’ and ‘stop’ trials were presented in a random order, which introduced temporal uncertainty. Therefore, this result is in accordance with previous work on temporal uncertainty difficulties in PWS and may be the result of an impairment in internal cueing.

An alternative explanation is that PWS show longer reaction times because they were in a state of heightened inhibition as the task demands required enacting a stopping response (at an unpredictable time) and might have prevented them from generating a ‘go’ response as quickly as PWTF. This would be consistent with predictions from the global suppression hypothesis (Neef et al., 2018, 2016). Nevertheless, the thresholding we implemented in the behavioural paradigm was adaptive so there was no benefit of being slower to perform the ‘go’ response to improve the success of stopping responses and participants were encouraged to perform the task as quickly as possible. Furthermore, the lack of difference in both the SSD and the SSRT during the stop trials suggests that inhibition acted to suppress go responses, rather than over-exerting stopping responses. Our behavioural results cannot distinguish between these two explanations (i.e. temporal uncertainty and global suppression).

### fMRI results

During ‘go’ responses, PWTF showed the expected focal activation of the left precentral gyrus (encompassing the hand area), left putamen extending to the opercular cortex, the SMA, and the cerebellum bilaterally, which is consistent with performing a button press with the right index finger. Compared with PWTF, PWS showed significant bilateral overactivation of areas comprising the typical movement ‘inhibition’ network but this was seen during ‘go’ trials. These areas include the inferior frontal gyrus, caudate nucleus and putamen bilaterally as well as the left precentral gyrus (encompassing the hand area). One explanation for these results (as for the behavioural data) is that PWS were consistently in a heightened inhibition state, i.e. areas of the inhibition network were more active, generally. Again, this interpretation is in accordance with predictions from the global response suppression hypothesis (Neef et al., 2018, 2015) and previous findings of hyperactivity in PWS in the basal ganglia, thalamus and substantia nigra during response preparation in a Go/NoGo task (Metzger et al., 2018). An alternative explanation is that the stuttering participants were in a higher state of arousal (possibly due to increased desire to perform the task well, or in response to being scanned). This arousal may cause general over activation of the areas involved in the task. However, this latter interpretation is unlikely to explain these results. There was not simply an amplification of all areas seen in PWTF; during ‘go’ trials, the right IFG was not active in PWTF but was active in PWS. The hypothesised importance of the right IFG in inhibition makes the former hypothesis more likely.

Our results indicate that PWTF activated the right IFG, frontal operculum and anterior insula during stopping responses, but not during ‘go’ responses, supporting the idea that this area is selective for stopping behaviour in the typical brain (Aron et al., 2007, 2003, 2004; Aron & Poldrack, 2006; Xue et al., 2008). PWS, on the other hand, activated right frontal regions (including IFG extending to the anterior insula) during both ‘go’ and ‘stop’ trials. Accordingly, during ‘go’ trials, PWS activated right IFG significantly more than PWTF. This indicates that the right IFG is active more generally in PWS, again in accord with the global suppression hypothesis (Metzger et al., 2018; Neef et al., 2018, 2011). However, it is important to note that although there were qualitative differences in the amount of activation during stopping behaviour between PWS and PWTF (see Figures 4 and 5), these differences failed to reach statistical significance at conventional thresholds (Z >3.1); to check for false negatives, we reduced the threshold to Z > 2.3 and found no group differences for the stop trials. Overall, this work supports the idea that PWS activate the right frontal regions irrespective of the type of response, but that the degree of activation during ‘stop’ trials was not statistically different to PWTF.

**Figure 4.**
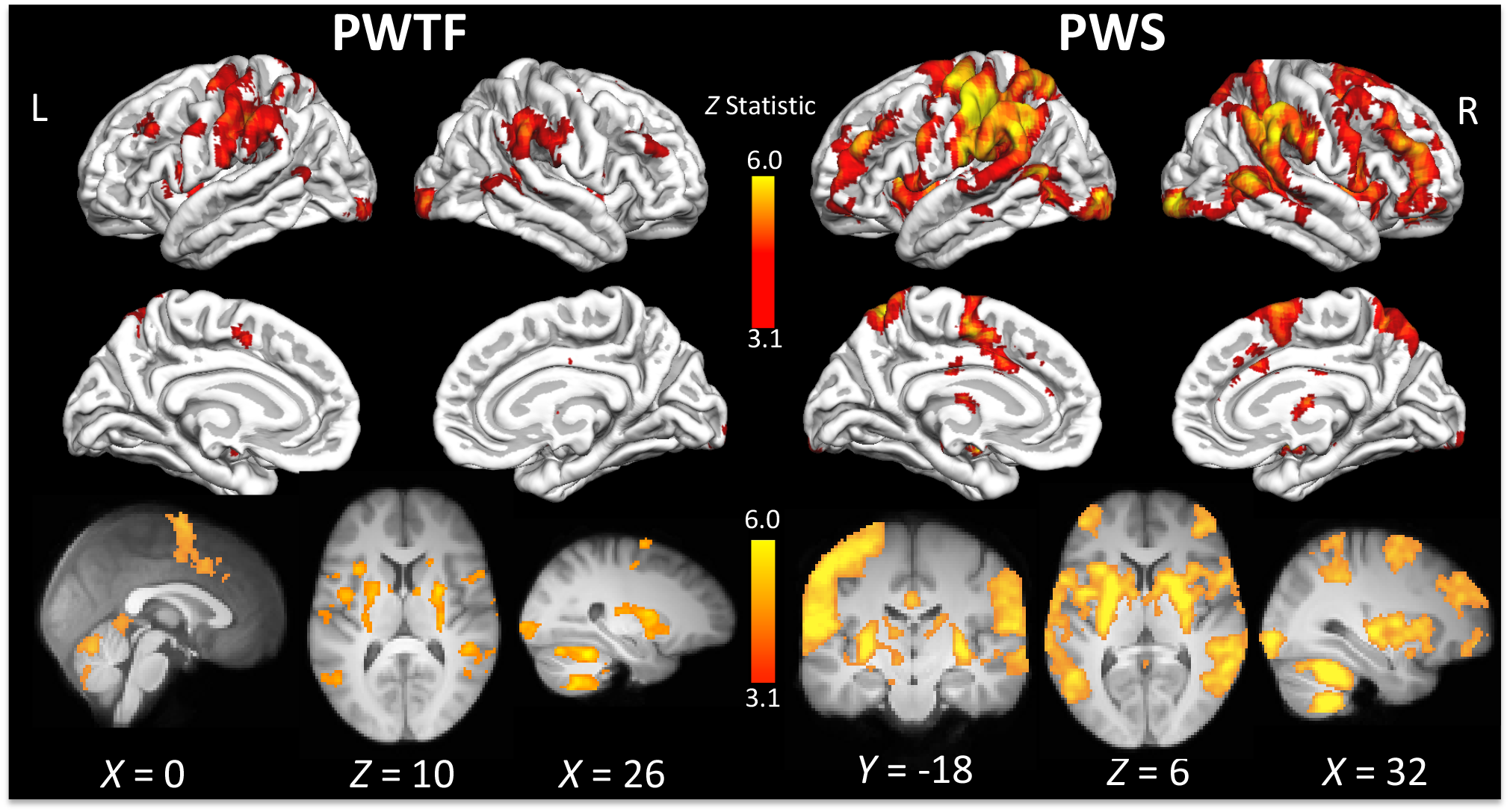
Activity during ‘successful stop’ trials. Coloured areas indicate statistical maps (thresholded at Z>3.1) overlaid on the cortical surface using FreeSurfer or on slices through the brain volume at the coordinate indicated below each image. L – left; R – right. See Tables 6 and 7 for a list of areas significantly activated for each group

**Figure 5.**
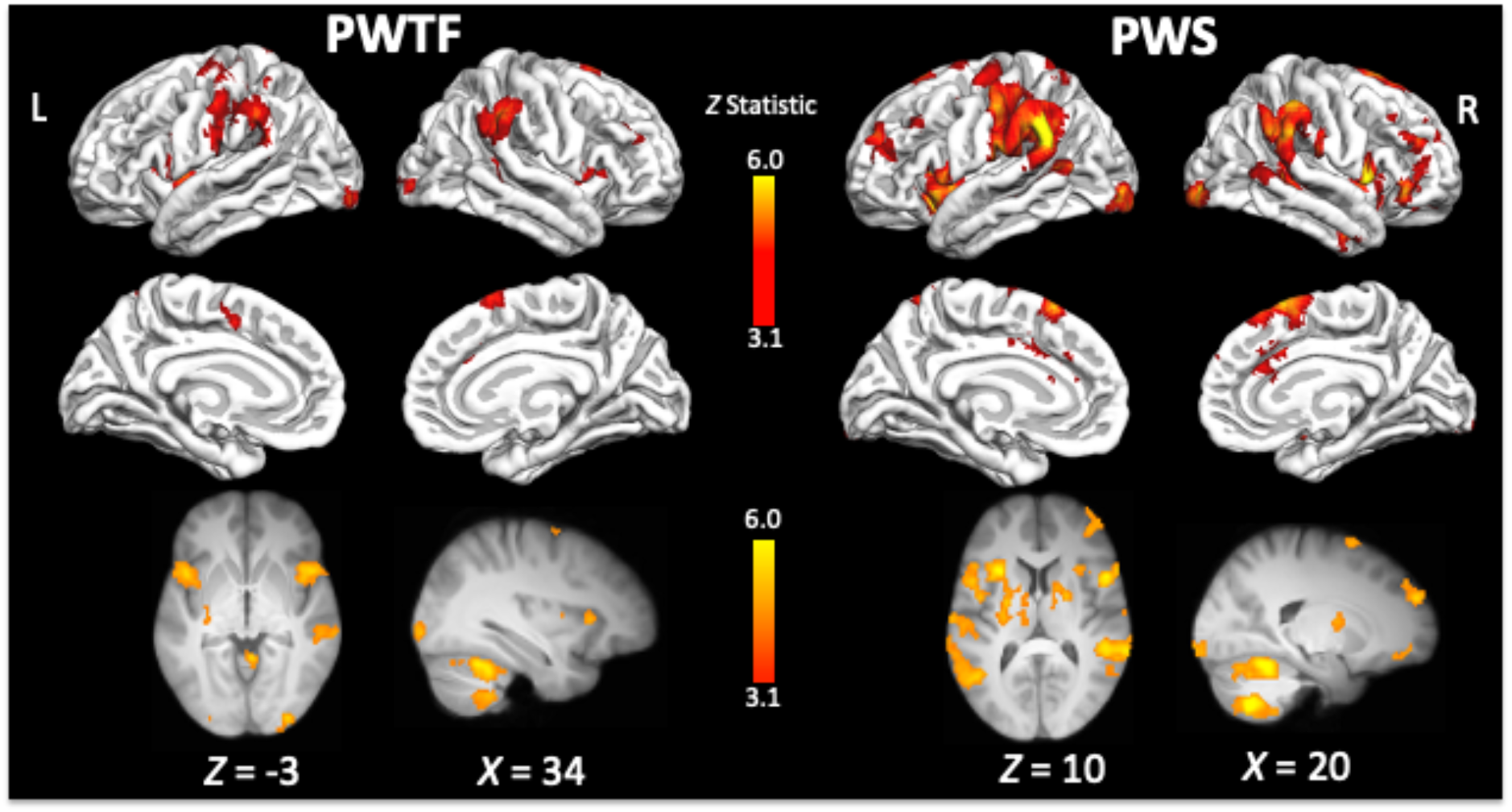
Activity during ‘unsuccessful stop’ trials. See legend to Figure 4 for details. See Tables 8 and 9 for a list of areas significantly activated for each group.

Whilst the right IFG has been a particular focus of the inhibition literature, it sits within a network of cortical-subcortical regions that carefully balance initiation and inhibition behaviour. During successful ‘stop’ trials, PWTF also activated the putamen, postcentral gyrus, supramarginal gyrus, and cerebellum bilaterally, and the SMA extending to the cingulate motor area. These areas have been implicated in previous studies of motor inhibition. Both direct and indirect pathways from these cortical areas via the putamen project back to the cortex via the thalamus to balance excitatory and inhibitory control (Alm, 2004; Burghaus et al., 2006; Giraud et al., 2008). The supramarginal gyrus was also activated in the inhibition of manual and spoken responses (Xue et al., 2008). Finally, the cingulate cortex is implicated in the cognitive control of inhibition. For example, the cingulate motor area was robustly activated (with a specific pattern for eye, hand or speech movement) when participants inhibited a congruent response in favour of a incongruent response (Paus, Petrides, Evans, & Meyer, 1993). These findings highlight the role of the cingulate cortex in the control of the balance between the selection of motor responses and active suppression of others (Paus, 2001; Paus et al., 1993).

The lack of statistical difference between the groups during ‘stop’ trials may reflect similar inhibitory control for stop trials in PWS and PWTF. This is against the prediction, based on the global suppression hypothesis, that PWS would show overactivity in key regions of the stopping network. However, visual inspection shows clear qualitative differences between the patterns of neural activation of stopping responses between PWS and PWTF, with PWS showing an amplified version of the controls. Therefore, while there are no differences that survive statistical thresholding, it cannot be ruled out that there are small differences between groups, or that other factors limit our ability to detect a significant difference between groups. One reason may relate to the design of the study. There were a large number of ‘go’ trials (144) but fewer stop trials (48) because the task requires stop trials to be unpredictable and in the minority. These stop trials are further divided into approximately 50% successful stops (~24) and 50% unsuccessful stops (~24). This reduced the power to detect differences between groups and may explain why we had sufficient power to detect group differences on the ‘go’ trials but not the ‘stop’ trials. Another factor is variability within the groups. Even though our sample of 31 was large for an imaging study, stuttering populations show considerable inter-individual differences (Yairi & Ambrose, 2013).

## Conclusions

We found that PWS were slower to respond to simple ‘go’ stimuli than PWTF, but there was no difference in stopping behaviour. Our fMRI results were consistent with these behavioural results. PWS showed significant overactivity of the inhibition network even during ‘go’ trials, which supports the idea of a global suppression mechanism in PWS. In addition, there were qualitative differences in the neural stopping response between groups, with PWS appearing to overactivate the inhibitory control network compared with PWTF. However, it must be stressed that these differences did not pass statistical significance, and that the study may have been underpowered to detect them. Overall, this study offers tentative support to the global suppression hypothesis of stuttering.

## Acknowledgements

We would like to thank all of our participants who took part in this study. We would also like to thank Professor Gui Xue (Beijing Normal University) for providing the code to run the stop signal task, Dr. Gabriel Cler for useful discussions, Louisa Needham for her assistance with data collection and members of the OHBA center MRI team (part of the WIN) Juliet Semple, Nicky Aikin, Sebastien Reiger, and Nicola Filippini.

Charlotte Wiltshire is supported by a DPhil scholarship from the Economic and Social Research Council UK [ES/J500112/1] and the Engineering and Physical Science Research Council UK [EP/N509711/1].

This work was supported by the Medical Research Council UK grant [MR/N025539/1]. The Wellcome Centre for Integrative Neuroimaging is supported by core funding from the Wellcome Trust [203139/Z/16/Z].

## References

Alm, P. A. (2004). Stuttering and the basal ganglia circuits: A critical review of possible relations. Journal of Communication Disorders, 37(4), 325–369. https://doi.org/10.1016/j.jcomdis.2004.03.001

Aron, A. R., Behrens, T. E., Smith, S., Frank, M. J., & Poldrack, R. A. (2007). Triangulating a cognitive control network using diffusion-weighted Magnetic Resonance Imaging (MRI) and functional MRI. Journal of Neuroscience, 27(14), 3743–3752. https://doi.org/10.1523/JNEUROSCI.0519-07.2007

Aron, A. R., Fletcher, P. C., Bullmore, E. T., Sahakian, B. J., & Robbins, T. W. (2003). Stop-signal inhibition disrupted by damage to right inferior frontal gyrus in humans. Nature Neuroscience, 6(2), 115–116. https://doi.org/10.1038/nn1003

Aron, A. R., & Poldrack, R. A. (2006). Cortical and subcortical contributions to stop signal response inhibition: Role of the subthalamic nucleus. Journal of Neuroscience, 26(9), 2424–2433. https://doi.org/10.1523/JNEUROSCI.4682-05.2006

Aron, A. R., Robbins, T. W., & Poldrack, R. A. (2004, April). Inhibition and the right inferior frontal cortex. Trends in Cognitive Sciences, Vol. 8, pp. 170–177. https://doi.org/10.1016/j.tics.2004.02.010

Belyk, M., Kraft, S. J., & Brown, S. (2017). Stuttering as a trait or a state revisited: motor system involvement in persistent developmental stuttering. European Journal of Neuroscience, 45(4), 622–624. https://doi.org/10.1111/ejn.13512

Brown, S., Ingham, R. J., Ingham, J. C., Laird, A. R., & Fox, P. T. (2005). Stuttered and fluent speech production: An ALE meta-analysis of functional neuroimaging studies. Human Brain Mapping, 25(1), 105–117. https://doi.org/10.1002/hbm.20140

Budde, K. S., Barron, D. S., & Fox, P. T. (2014). Stuttering, induced fluency, and natural fluency: A hierarchical series of activation likelihood estimation meta-analyses. Brain and Language, 139, 99–107. https://doi.org/10.1016/j.bandl.2014.10.002

Burghaus, L., Hilker, R., Thiel, A., Galldiks, N., Lehnhardt, F. G., Zaro-Weber, O., … Heiss, W. D. (2006). Deep brain stimulation of the subthalamic nucleus reversibly deteriorates stuttering in advanced Parkinson’s disease. Journal of Neural Transmission, 113(5), 625–631. https://doi.org/10.1007/s00702-005-0341-1

Chambers, C. D., Bellgrove, M. A., Gould, I. C., English, T., Garavan, H., McNaught, E., … Mattingley, J. B. (2007). Dissociable mechanisms of cognitive control in prefrontal and premotor cortex. Journal of Neurophysiology, 98(6), 3638–3647. https://doi.org/10.1152/jn.00685.2007

Chen, W., De Hemptinne, C., Miller, A. M., Lim, D. A., Larson, P. S., Starr, P. A., … Lim, D. A. (2020). Prefrontal-subthalamic hyperdirect pathway modulates movement inhibition in humans. Neuron, 0(0), 1–10. https://doi.org/10.1016/j.neuron.2020.02.012

Chevrier, A. D., Noseworthy, M. D., & Schachar, R. (2007). Dissociation of response inhibition and performance monitoring in the stop signal task using event-related fMRI. Human Brain Mapping, 28(12), 1347–1358. https://doi.org/10.1002/hbm.20355

De Nil, L., Kroll, R. M., Lafaille, S. J., & Houle, S. (2003). A positron emission tomography study of short- and long-term treatment effects on functional brain activation in adults who stutter. Journal of Fluency Disorders, 28(4), 357–380. https://doi.org/10.1016/j.jfludis.2003.07.002

Eggers, K., De Nil, L., & Van Der Bergh, B. R. H. (2013). Inhibitory control in childhood stuttering. Journal of Fluency Disorders, 38(1), 1–13. https://doi.org/10.1016/j.jfludis.2012.10.001

Fox, P. T., Ingham, R. J., Ingham, J. C., Hirsch, T. B., Downs, J. H., Martin, C., … Lancaster, J. L. (1996). A PET study of the neural systems of stuttering. Nature, 382(6587), 158–162. https://doi.org/10.1038/382158a0

Giraud, A. L., Neumann, K., Bachoud-Levi, A. C., von Gudenberg, A. W., Euler, H. A., Lanfermann, H., & Preibisch, C. (2008). Severity of dysfluency correlates with basal ganglia activity in persistent developmental stuttering. Brain and Language, 104(2), 190–199. https://doi.org/10.1016/j.bandl.2007.04.005

Hartwigsen, G., Neef, N. E., Camilleri, J. A., Margulies, D. S., & Eickhoff, S. B. (2019). Functional Segregation of the Right Inferior Frontal Gyrus: Evidence from Coactivation-Based Parcellation. Cerebral Cortex, 29(4), 1532–1546. https://doi.org/10.1093/cercor/bhy049

Heuer, R. J., Sataloff, R. T., Mandel, S., & Travers, N. (1996). Neurogenic stuttering: Further corroboration of site of lesion. Ear, Nose and Throat Journal, 75(3), 161–168. https://doi.org/10.1177/014556139607500312

Kell, C. A., Neumann, K., von Kriegstein, K., Posenenske, C., von Gudenberg, A. W., Euler, H. A., & Giraud, A. L. (2009). How the brain repairs stuttering. Brain, 132(10), 2747–2760. https://doi.org/10.1093/brain/awp185

Markett, S., Bleek, B., Reuter, M., Prüss, H., Richardt, K., Müller, T., … Montag, C. (2016). Impaired motor inhibition in adults who stutter ‒evidence from speech-free stop-signal reaction time tasks. Neuropsychologia, 91, 444–450. https://doi.org/10.1016/j.neuropsychologia.2016.09.008

Metzger, F. L., Auer, T., Helms, G., Paulus, W., Frahm, J., Sommer, M., & Neef, N. E. (2018). Shifted dynamic interactions between subcortical nuclei and inferior frontal gyri during response preparation in persistent developmental stuttering. Brain Structure and Function, 223(1), 165–182. https://doi.org/10.1007/s00429-017-1476-1

Mink, J. W. (1996). The basal ganglia: Focused selection and inhibition of competing motor programs. Progress in Neurobiology, 50(4), 381–425. https://doi.org/10.1016/S0301-0082(96)00042-1

Nambu, A., Tokuno, H., & Takada, M. (2002). Functional significance of the cortico-subthalamo-pallidal “hyperdirect” pathway. Neuroscience Research, 43(2), 111–117. https://doi.org/10.1016/S0168-0102(02)00027-5

Neef, N. E., Anwander, A., Bütfering, C., Schmidt-Samoa, C., Friederici, A. D., Paulus, W., & Sommer, M. (2018). Structural connectivity of right frontal hyperactive areas scales with stuttering severity. Brain, 141(1), 191–204. https://doi.org/10.1093/brain/awx316

Neef, N. E., Anwander, A., & Friederici, A. D. (2015). The Neurobiological Grounding of Persistent Stuttering: from Structure to Function. Current Neurology and Neuroscience Reports, Vol. 15. https://doi.org/10.1007/s11910-015-0579-4

Neef, N. E., Bütfering, C., Anwander, A., Friederici, A. D., Paulus, W., & Sommer, M. (2016). Left posterior-dorsal area 44 couples with parietal areas to promote speech fluency, while right area 44 activity promotes the stopping of motor responses. NeuroImage, 142, 628–644. https://doi.org/10.1016/j.neuroimage.2016.08.030

Neef, N. E., Jung, K., Rothkegel, H., Pollok, B., von Gudenberg, A. W., Paulus, W., & Sommer, M. (2011). Right-shift for non-speech motor processing in adults who stutter. Cortex, 47(8), 945–954. https://doi.org/10.1016/j.cortex.2010.06.007

Paus, T. (2001, June). Primate anterior cingulate cortex: Where motor control, drive and cognition interface. Nature Reviews Neuroscience, Vol. 2, pp. 417–424. https://doi.org/10.1038/35077500

Paus, T., Petrides, M., Evans, A. C., & Meyer, E. (1993). Role of the human anterior cingulate cortex in the control of oculomotor, manual, and speech responses: A positron emission tomography study. Journal of Neurophysiology, 70(2), 453–469. https://doi.org/10.1152/jn.1993.70.2.453

R Core Team. (2019). R: A language and environment for statistical computing. R Foundation for Statistical Computing.

Ray Li, C. S., Yan, P., Sinha, R., & Lee, T. W. (2008). Subcortical processes of motor response inhibition during a stop signal task. NeuroImage, 41(4), 1352–1363. https://doi.org/10.1016/j.neuroimage.2008.04.023

Tourville, J. A., & Guenther, F. H. (2010). Language and Cognitive Processes The DIVA model: A neural theory of speech acquisition and production. https://doi.org/10.1080/01690960903498424

Tourville, J. A., Reilly, K. J., & Guenther, F. H. (2008). Neural mechanisms underlying auditory feedback control of speech. NeuroImage, 39(3), 1429–1443. https://doi.org/10.1016/j.neuroimage.2007.09.054

Van Borsel, J., Van Der Made, S., & Santens, P. (2003). Thalamic stuttering: A distinct clinical entity? Brain and Language, 85(2), 185–189. https://doi.org/10.1016/S0093-934X(03)00061-0

Watkins, K. E., Smith, S., Davis, S., & Howell, P. (2008). Structural and functional abnormalities of the motor system in developmental stuttering. Brain, 131(1), 50–59. https://doi.org/10.1093/brain/awm241

Wu, J. C., Maguire, G., Riley, G., Lee, A., Keator, D., Tang, C., … Najafi, A. (1997). Increased dopamine activity associated with stuttering. NeuroReport, 8(3), 767–770. https://doi.org/10.1097/00001756-199702100-00037

Xue, G., Aron, A. R., & Poldrack, R. A. (2008). Common neural substrates for inhibition of spoken and manual responses. Cerebral Cortex, 18(8), 1923–1932. https://doi.org/10.1093/cercor/bhm220

Yairi, E., & Ambrose, N. G. (2013). Epidemiology of stuttering: 21st century advances. Journal of Fluency Disorders, 38(2), 66–87. https://doi.org/10.1016/j.jfludis.2012.11.002

